# Flex-sweep 2.0: more flexible and faster selective sweeps detection

**DOI:** 10.64898/2026.08.06.743046

**Authors:** Jesús Murga-Moreno, David Enard

## Abstract

Flex-sweep is a convolutional neural network-based method able to detect a wide range of selective sweeps, including those thousands of generations old, from single population genomic data, while robust to background selection. Here we present a substantial update that streamlines the entire workflow. The new version vastly reduces memory needs and vastly speeds up summary-statistic computation over fully customizable statistics combinations and genomic regions, relaxes CNN constraints by supporting custom architectures and haplotype matrix sorting methods. Domain-Adaptive Neural Network (DANN) training is now supported, as well as ancestral-state polarization and a robust, clustering and confounder-aware gene set sweep enrichment pipeline robust for downstream analysis. Flex-sweep 2.0 scales to hundreds of thousands of training simulations, and enables genome-wide inference on a standard workstation.

## Introduction

Quantifying the contribution of selective sweeps to genomic adaptation remains an active area of research in population genetics, and developing methods able to detect diverse sweeps remains a challenge. Machine Learning (ML) techniques have provided robust and versatile alternatives to classical approaches based on summary statistics, [Kim and Nielsen, 2004, Voight et al., 2006, Garud et al., 2015, Akbari et al., 2018], maximum likelihood [Williamson et al., 2007, DeGiorgio et al., 2016, Harris and DeGiorgio, 2020], composite multiple signals [Grossman et al., 2010, Sugden et al., 2018, Alachiotis and Pavlidis, 2018] or Approximate Bayesian Computation (ABC) [Peter et al., 2012, Racimo et al., 2014, Johri et al., 2023]. ML-based approaches (SVM, RNN, CNN, GNN, DANN among others [Pavlidis et al., 2010, Kern and Schrider, 2018, Hejase et al., 2022, Mo and Siepel, 2023, van den Belt and Alachiotis, 2025, Whitehouse et al., 2024]) can learn intrinsic patterns in abstract haplotype images [Arnab et al., 2025, van den Belt and Alachiotis, 2025, Zhao and Alachiotis, 2025] or feature vectors of classical summary statistics [Kern and Schrider, 2018, Caldas et al., 2022, Lauterbur et al., 2023], typically acting as image classifiers trained on neutral and sweep segments to then classify empirical genomic windows.

Most existing methods have been limited to specific sweep types, ages or starting and ending frequencies. Detecting older sweeps is especially challenging [Lauterbur et al., 2023]. Methods aimed at detecting old sweeps rely on allele-frequency differences between populations, (i) limiting scans to populations with outgroup and/or population data [Racimo et al., 2014, Key et al., 2016, Racimo, 2016, Cheng et al., 2017, Peyrégne et al., 2017] and (ii) creating vulnerability to background selection [Hernandez et al., 2011].

While Flex-sweep is able to detect diverse sweeps across a wide range of ages from a single phased population, all the while being robust to background selection [Lauterbur et al., 2023], we and other users have found that the first version has important limitations, mainly a heavy computation burden and sensitivity of the results to differences between simulated and tested recombination rates (see below). Here we present a faster, more flexible update that streamlines the entire pipeline, making Flex-sweep a general-purpose tool for genome-wide selection scans. We also included new features to facilitate non-model species analysis such as robustness to mis-specification through Domain-Adaptive Neural Network (DANN) training and ancestral-allele polarization, as well as customizable statistics and region feature-vector computation and haplotype-matrix support.

### Pipeline and new features

To improve efficiency, robustness and reproducibility, we refactored Flex-Sweep as a user-friendly, well-documented Python package with a Command Line Interface, avoiding non-standard input files and manual installation of external dependencies.

We tested Flex-sweep 2.0 on larger sets of training simulations, predicted 1000GP human populations [Byrska-Bishop et al., 2022], and benchmarked timing and scalability against a widely used summary-statistic CNN Kern and Schrider [2018] (see Computational Resources). Similarly to the first version, Flexsweep works in three main steps: simulation, summary statistics estimation (feature vectors), and training/classification.

#### Simulations

The simulations module now takes advantage of demes [Gower et al., 2022] to simulate custom demography histories, and the software automatically compiles discoal binary [Kern and Schrider, 2016], to avoid external installation. Priors parameters are flexible to different distribution configurations and do not require manually fitting a custom file, but any option is documented and accessible through the Python API or CLI.

#### Feature vector estimation

We refactored the entire feature-vector module for speed and traceability using numpy and scikit-allel data structures. The statistics were reimplemented through numba [Lam et al., 2015] and numpy vectorization except for *nS*_*L*_ and *iHS*, which use scikit-allel functions. Beyond the statistics first introduced in Flexsweep 1.0 (*Sratio, hapDAF* -*o*/*s, highfreq, lowfreq*) [Lauterbur et al., 2023], we also refactored *iSAFE* [Akbari et al., 2018], *DIND* [Barreiro et al., 2009], *and the modified H*12 [Garud et al., 2015] and *HAF* [Ronen et al., 2015] (see Supplementary material).

Outputs and feature vectors are stored in numpy arrays and polars DataFrames [Vink et al., 2026], avoiding intermediate files, custom project structure and high storage requirements of the previous design while minimizing RAM and keeping the pipeline fully traceable. The new design allows us to expose raw SNP and window-level information for complementary analysis and provides an entire new module for classical outlier scans over empirical distributions [Akey, 2009].

Importantly, users can now combine any included summary statistic (see Table 1) over user-defined genomic intervals to fit the most informative statistic and region combination for a given organism. We also include two widely used balancing-selection statistics, providing a potential interface for balancing-selection analyses (see Isildak et al. [2021] for further discussion).

**Table 1:** Summary statistics implemented in Flex-sweep.

| Summary statistic | Description | Summarizes | Citation |
| --- | --- | --- | --- |
| $\pi$ | Average number of pairwise nucleotide differences | Nucleotide diversity | Tajima [1983] |
| $\theta_w$ | Watterson’s estimator based on segregating sites | Nucleotide diversity | Watterson [1975] |
| Fu & Li’s $D$ and $D^*$ | Excess of singletons relative to total mutations; $D$ (polarized, requires outgroup), $D^*$ (folded) | SFS | Fu and Li [1993] |
| Fu & Li’s $F$ and $F^*$ | Excess of singletons relative to average pairwise differences; $F$ (polarized, requires outgroup), $F^*$ (folded) | SFS | Fu and Li [1993] |
| $\theta_h$ | Excess of high-frequency derived alleles | SFS | Fay and Wu [2000] |
| Fay & Wu’s $H$ and $H'$ | Excess of high-frequency derived alleles | SFS | Fay and Wu [2000], Zeng et al. [2006] |
| Zeng’s $E$ | Contrast between $\theta_\pi$ and Fay & Wu’s $H$ | SFS | Zeng et al. [2006] |
| Achaz’s $Y$ and $Y^*$ | Tajima’s $D$ analogue excluding singleton sites, robust to sequencing errors; $Y$ excludes derived singletons (polarized, requires outgroup), $Y^*$ excludes minor-allele singletons (folded) | SFS | Achaz [2008] |
| Achaz’s $T_\Omega$ | Generalised neutrality test statistic defined as the normalized difference between any two frequency-spectrum-based $\hat{\theta}$ estimators, under a unified framework that expresses all classical estimators and tests as linear combinations of the site frequency spectrum | SFS | Achaz [2009] |
| *hapDAF-o* | Haplotype-derived allele frequency (old) | SFS | Lauterbur et al. [2023] |
| *hapDAF-s* | Haplotype-derived allele frequency (standing) | SFS | Lauterbur et al. [2023] |
| *DIND* | Derived intra-allelic nucleotide diversity | Diversity on derived background | Barreiro et al. [2009] |
| *Sratio* | Segregating sites ratio | Diversity on derived background | Lauterbur et al. [2023] |
| *lowfreq* | Low-frequency alleles on derived background | Diversity on derived background | Lauterbur et al. [2023] |
| *highfreq* | High-frequency alleles on derived background | Diversity on derived background | Lauterbur et al. [2023] |
| *iHS* (scikit-allele) | Integrated haplotype score | Haplotype structure | Voight et al. [2006] |
| Δ-*iHH* (scikit-allele) | Absolute iHH difference between ancestral and derived alleles | Haplotype structure | Grossman et al. [2010] |
| $nS_L$ (scikit-allele) | Number of segregating sites by length | Haplotype structure | Ferrer-Admetlla et al. [2014] |
| Garud’s $H$ | Haplotype homozygosity statistics (including $H1$, $H12$, and $H2/H1$) | Haplotype structure | Garud et al. [2015] |
| *iSAFE* | Integrated selection of allele favored by evolution | Haplotype structure | Akbari et al. [2018] |
| *HAF* | Haplotype allele frequency | Haplotype structure | Ronen et al. [2015] |
| h-scan | Average pairwise haplotype homozygosity tract length | Haplotype structure | Schlamp et al. [2016] |
| Modified $H12$ | Frequencies of first and second most common haplotypes, modified to use 80% identity threshold | Haplotype structure | Lauterbur et al. [2023] |
| Haplotypes count | Distribution of differences between haplotypes | Haplotype structure | Kern and Schrider [2018] |
| $\text{Var}(d_{ij})$ | Variance of the distribution of pairwise haplotype mismatch distances $d_{ij}$ within a subwindow | Haplotype structure | Kern and Schrider [2018] |
| $\text{Skew}(d_{ij})$ | Skewness of the distribution of pairwise haplotype mismatch distances $d_{ij}$ within a subwindow | Haplotype structure | Kern and Schrider [2018] |
| $\text{Kurt}(d_{ij})$ | Excess kurtosis of the distribution of pairwise haplotype mismatch distances $d_{ij}$ within a subwindow | Haplotype structure | Kern and Schrider [2018] |
| $\text{DAF}_{\max}$ | Maximum derived allele frequency in a subwindow | Site frequency spectrum | Kern and Schrider [2018] |
| Kelly’s $Z_{nS}$ | Average linkage disequilibrium ($r^2$) between segregating sites | Linkage disequilibrium | Kelly [1997] |
| $\omega_{\max}$ | Maximum LD between selected and flanking regions | Linkage disequilibrium | Kim and Nielsen [2004] |
| LASSI $T$ and $\hat{m}$ | Likelihood-based detection of selective sweeps using haplotype structure | Haplotype structure | Harris and DeGiorgio [2020] |
| RAiSD $\mu$ | Composite detection of selective sweeps using SFS, LD, and diversity | Composite likelihood | Alachiotis and Pavlidis [2018] |
| $\beta_{(std)}^{(1)*}$ | Correlation of allele frequency with local polymorphism | SFS (balancing selection) | Siewert and Voight [2020] |
| NCD1 | Non-central deviation of the SFS from neutrality | SFS (balancing selection) | Bitarello et al. [2018] |

#### Normalization

Recombination rate heterogeneity biases population genetic inferences broadly, including ancestral recombination graph inference and effective population size estimation [Booker et al., 2020, Johnson and Voight, 2018, Ishigohoka and Liedvogel, 2025]. In the context of sweep detection, classical summary statistics show heterogeneous distributions under different recombination rates even under strict neutrality, which can increase false positive rates (FPR) [Johnson and Voight, 2018, Booker et al., 2020]. Johnson and Voight [2018] showed that the spurious correlation of haplotype-based statistics with local recombination rate can be addressed through joint stratification by allele frequency and recombination rate. Other methods may be less exposed to this effect, depending on the statistics they use and on their own normalization strategies. Inspecting the distribution of each statistic across recombination rate bins, however, we found that the statistics specifically developed for Flex-sweep are also sensitive to local recombination rate, while contributing substantially to classification [Lauterbur et al., 2023] (see Supplementary Figure S1). Flex-sweep therefore incorporates recombination-rate-stratified normalization following Johnson and Voight [2018], for both SNP-centered and window-centered statistics, whenever a recombination map is available. This removes the variation attributable to recombination heterogeneity while preserving the discriminative information (see Supplementary Figure S2), and makes the classifier more robust to discrepancies between training and test sets.

For window-centered statistics, each input is standardized within bins defined by the joint combination of window size, window center, and local recombination rate:

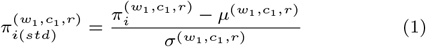

where *π*_*i*_ is a given statistic, and 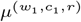 and 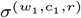 are the mean and standard deviation computed across all windows sharing the same window size *w*_1_, center *c*_1_, and recombination rate bin *r* ∈ {1, …, *R*}, with bin edges defined from the empirical recombination rate distribution. For SNP-based statistics, normalization is analogous but also conditional on derived allele frequency (DAF):

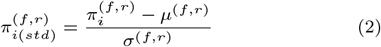

where *f* denotes the DAF bin of SNP *i*, and *µ*^(*f,r*)^ and *σ*^(*f,r*)^ are the mean and standard deviation computed across all SNPs falling within the same DAF bin *f* and recombination rate bin *r*.

In both cases the recombination rate bin edges are defined from the empirical recombination rate distribution. For the human data analysed here (see section Application to human data), we derived them from the deCODE recombination map [Halldorsson et al., 2019], interpolating genome-wide 1.2 Mb regions to obtain 10 bins with upper edges [0.37, 0.55, 0.71, 0.87, 1.05, 1.26, 1.53, 1.88, 2.46, 6.1] cM/Mb, the first bin being unbounded below (regions below 0.01 cM/Mb were filtered before binning).

#### Training and prediction

Because feature vectors are now flexible to any combination of statistics, genomic centers, and window sizes, Flex-sweep will adapt to custom CNN architectures, supporting both 1D and 2D convolutions. By default the feature vectors are split into 80% training, 10% validation and 10% test sets, with early stopping monitoring the validation AUC. To further enable compatibility with haplotype image-based approaches, we incorporated a comprehensive set of sorting and rearrangement strategies prior to model training [Zhao and Alachiotis, 2025, Tran et al., 2025]. These include both column- and row-level sorting by derived allele frequency, correlation coefficients, occurrence frequency, and subregional bipartite correlation [Zhao and Alachiotis, 2025], as well as disruption-based permutations targeting linkage disequilibrium (LD), site frequency spectrum (SFS), and allele frequency patterns [Tran et al., 2025].

#### DANN

Flex-sweep is now more versatile to analyze non-model organisms when the availability of mutation rate, recombination rate, and demography estimates is limited. We therefore implemented the Domain Adaptive model proposed by Mo and Siepel [2023] to address discrepancies between training and tested sets. Domain Adaptive Neural Network (DANN) trains not only using labeled simulated data as expected for a CNN, but also incorporates empirical unlabeled data during the training. The goal is then to generalize the classification task across any feature distorting simulated feature vector distributions from real data by learning a shared representation pattern that is highly predictive for the CNN classifier but uninformative about the domain, i.e. whether the feature vector comes source (simulated domain) or target (empirical domain) data. In the case of ML approaches like CNN, trained models under unrealistic demography, recombination, or mutation landscape can easily confound sweep prediction due to the overfitting of artifacts [Mo and Siepel, 2023]. The DANN model is explicitly designed to account for and mitigate such a mismatch between simulated and real data. Note that when working with extremely out-of-range demographies or other simulated parameters, DANN implementation may still perform worse than the original CNN [Mo and Siepel, 2023].

To improve robustness under domain shift, the systematic difference between simulated and empirical feature distributions, we stabilize adversarial adaptation through controlled gradient scheduling. Rather than a fixed gradient-reversal factor, a time-epoch–dependent schedule ramps the adversarial signal from zero to a predefined maximum, making the model first learn task-discriminative features from labeled simulations before enforcing domain invariance. This way we prevent the domain discriminator from distorting the feature extractor early in training, when simulation-specific artifacts would otherwise dominate the gradient (see Supplementary Material)

#### Polarization

Given the impact of ancestral allele mis-specification on positive selection tests [Hernandez et al., 2007], Flex-sweep refactors and extends est-sfs [Keightley and Jackson, 2018] to support straightforward allele polarization, automatically retrieving multiple outgroup sequences from Multi-Alignment Format (MAF) files to infer the ancestral allele. Given a focal-species polymorphism dataset (VCF) and MAF file, we adapted maftk [Buffalo, 2025] parsing layer to i) subsets up to three outgroup species from the alignment ii) extracts the aligned outgroup bases at polymorphic sites, and iii) computes posterior probabilities per-site ancestral-state following Keightley and Jackson [2018]. These posteriors are then used to probabilistically assign ancestral and derived states for both major and minor alleles. We aim to exploit the advent of large comparative genomic resources—such as the Zoonomia Project [Armstrong et al., 2020, Kuderna et al., 2024] and the ability to incorporate additional species into whole-genome alignments [Armstrong et al., 2020] to enable curated selection scans in non-model species.

#### Rank and enrichment

We included a rank algorithm for post-processing sweep probabilities and associated genomic elements. The function relies on polars-bio [Wiewiórka et al., 2025] to quickly estimate genomic distances between the predicted genomic region and the element listed in a BED file to associate the nearest sweep prediction. To assess whether highly ranked elements are enriched within predefined genomic categories, we implemented the enrichment pipeline described in Enard and Petrov [2020], Di et al. [2021], which tests for significant enrichment among top-ranked genome-wide sweeps.

Briefly (see the two previous cited articles for more details), enrichment is evaluated across multiple rank thresholds by comparing tested gene sets of interest for each threshold to matched control sets, generated via an iterative bootstrap that samples controls genes accounting for confounding factors while controlling for distance and tolerance range. At each threshold, enrichment is summarized as the ratio of set of interest to mean control counts, with confidence intervals from the control-set distribution. This gene set enrichment pipeline also includes a step to estimate unbiased False Positive Rates (FPR) [Enard and Petrov, 2020, Di et al., 2021].

### Application to human data

Flex-sweep now scales to the entire 1000GP dataset [Byrska-Bishop et al., 2022], with the ability to handle larger training sets. We simulated 250,000 neutral and non-neutral 1.2Mb regions accounting for demography (versus 22,000 originally), as well as a for a wide variety of sweep signals (simulations prior parameters are described at Supplementary Material).

Because of the performance improvements we were able to simulate and train the 26 1000GP populations; similarly to Lauterbur et al. [2023] we present here the YRI population to easily compare results. Figure 1 shows Manhattan plot for YRI population genome-wide sweep predictions using the stratified recombination normalization previously described. The YRI baseline CNN reached ROC-AUC and PR-AUC ≈ 0.98, correctly classifying 96% of neutral and 91% of sweep regions at a 3.8% false positive rate (FPR) and 8.7% false negative rate (FNR) (Supplementary Figures S3-S4).

**Fig. 1:**
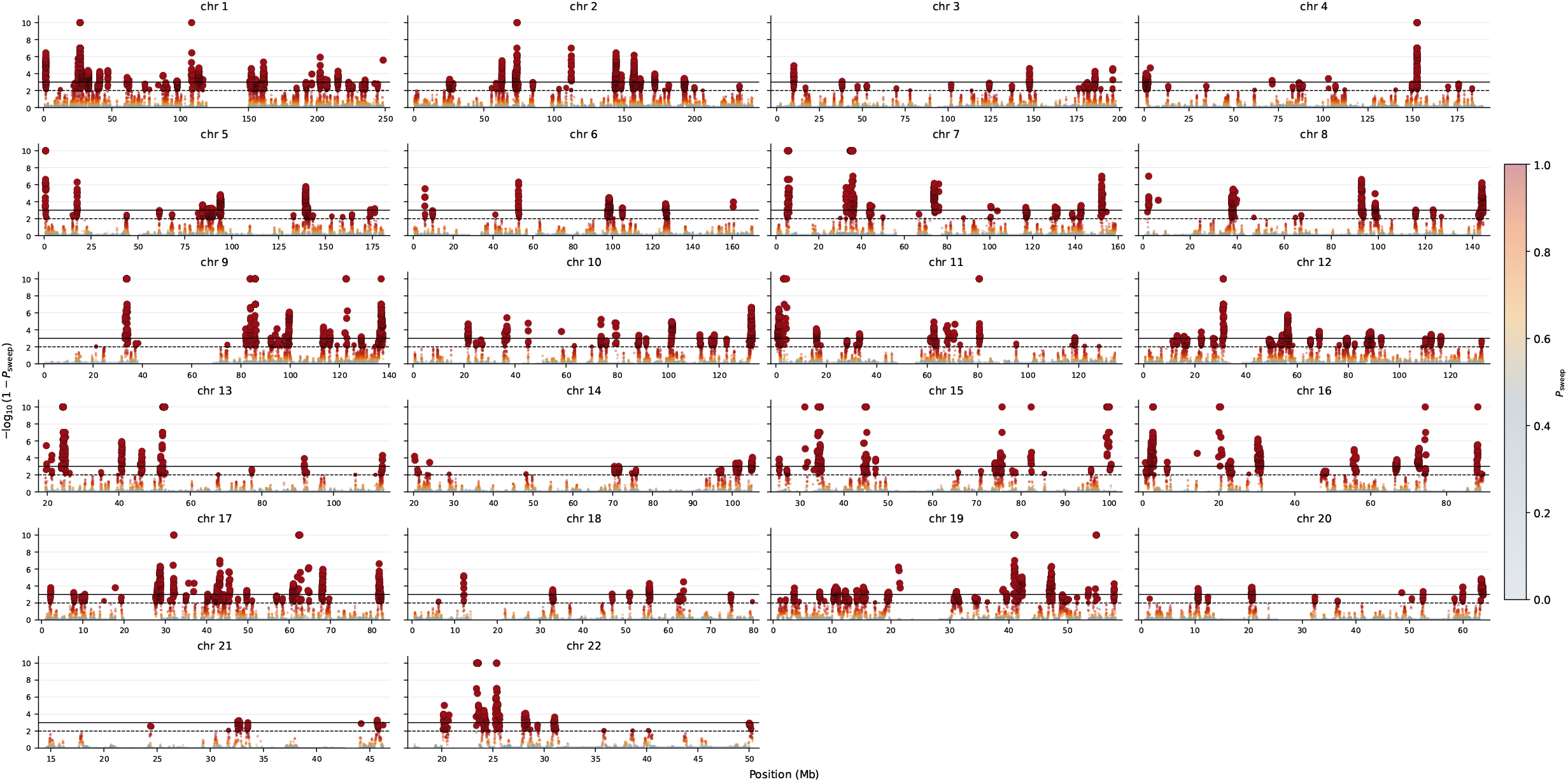
YRI predictions. Manhattan plot genome-wide sweep predictions over YRI population autosomes.

In addition, similarly to Lauterbur et al. [2023] we tested for both older strong hard sweep and background selection following Schrider [2020] model (see Figure S5), providing the baseline set up to test on real human data, where the action and background selection and potential sweep candidates due to ancient epidemics is pervasive [Lauterbur et al., 2023, Souilmi et al., 2021, Enard and Petrov, 2020]. For both cases we simulated 1,000 neutral and non-neutral regions setting up the same previously described mutation and recombination rate (see Supplementary Material). For the old hard sweep scenario the baseline CNN reached a ROC-AUC of 0.98, correctly classifying 91% of neutral and 96% of sweep regions at an 8.7% FPR and 4% FNR (Figure 2 and Supplementary Figure S6). Labeling both neutral and BGS regions as neutral, the CNN assigned sweep probabilities to BGS regions that were nearly indistinguishable from neutral ones, reaching a ROC-AUC of 0.598 (Figure 3) and less than 1% of BGS regions were classified as sweeps (Supplementary Figure S7), confirming that the classifier remains robust to BGS.

**Fig. 2:**
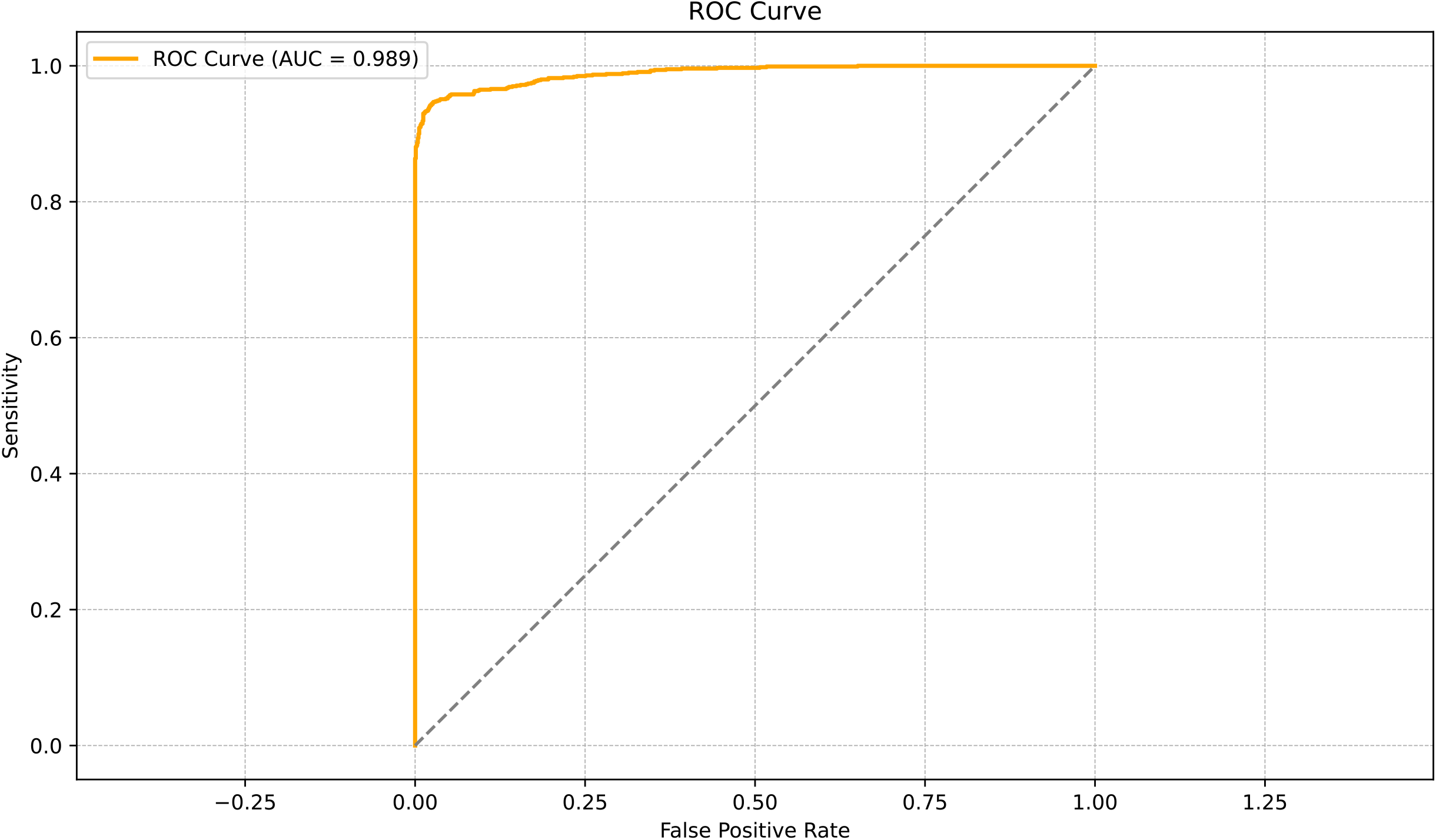
ROC curve prediction for old hard sweeps. 1,000 neutral and non-neutral simulations following YRI-like demography were tested accounting for timing distribution *t U* (5, 000, 10, 000). The prediction was performed using the baseline trained CNN accounting for 250,000 neutral and non-neutral regions with timing distribution of *t U* (0, 5000).

**Fig. 3:**
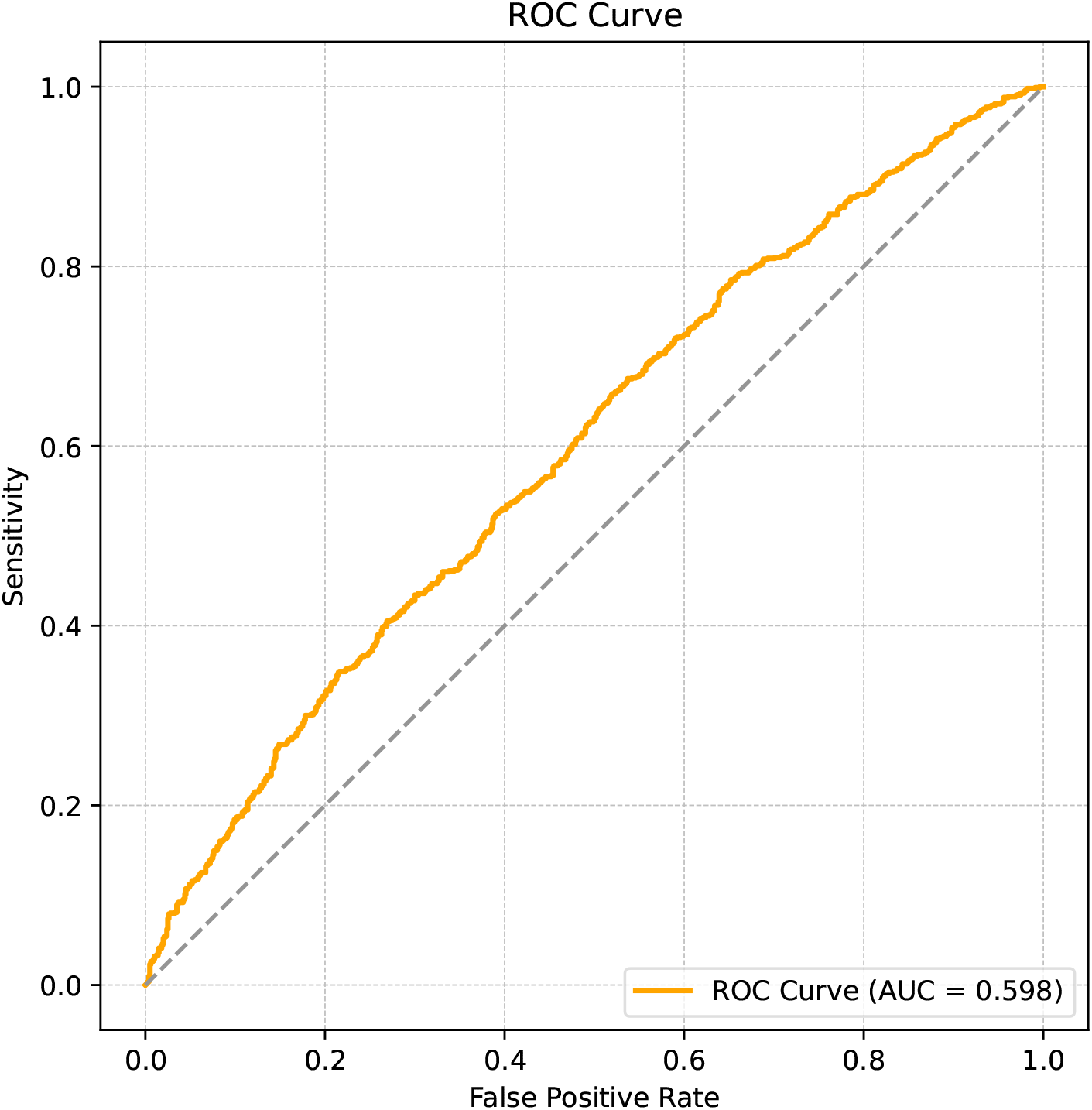
Roc curve for background selection (BGS) simulations. 1,000 neutral and non-neutral simulations were performed using stdpopsim under the Tennesen model. BGS were simulated under the Schrider and Kern [2017] central model following Kim et al. [2017] DFE. To test the amount of False Positive we labeled BGS simulations as sweeps.

### Computational Resources and Requirements

We developed Flex-sweep 2.0 using an AMD Ryzen 9 PRO 5945 12-core CPU and 64 GB of RAM, and tested thread scalability across the 26 1000GP populations on 2 AMD EPYC 7713 64-Core CPUs and 4 TB of RAM.

We compared against diploS/HIC [Kern and Schrider, 2018], a widely used CNN-based method (accessed 12/03/26), modifying its haploid mode to parallelize as in Flex-sweep, matching phasing, missing-data, and simulation-length assumptions. Across thread configurations and 22,000 neutral and non-neutral YRI-like simulations, Flex-sweep showed a 2–5-fold speed-up (Supplementary Figure S8). Genome-wide YRI computation showed similar speed-ups but little thread scaling, likely due to efficient statistic estimation, fewer analyzed windows (only unique center/window combinations estimated), and scheduler overhead. Normalization differences between methods should be negligible relative to statistic estimation and I/O time.

Training and prediction ran on an RTX 3080 with 10 GB of VRAM. YRI training took ≈ 20 minutes using 250,000 neutral and non-neutral feature vectors respectively, with early stopping and converging in about 10 epochs.

## Data availability

Flex-sweep source code is available at https://github.com/jmurga/flexsweep and the manual can be found in https://flexsweep.readthedocs.io/en/latest/. Flex-sweep can be easily installed through the PyPi.

## Author contributions

J.M-M. and D.E. designed the study, and wrote the manuscript. J.M-M implemented Flex-sweep v2.0. J.M-M and D.E. analyzed the data.

## Acknowledgment

We thank Andrew Kern for helpful discussion about discoal RAM usage and the new discoal implementation.

## Supplementary material

### Simulations

We used discoal [Kern and Schrider, 2016] to perform 250,000 neutral and non-neutral 1.2Mb regions for each 1000GP population, following ancestral population sizes retrieved from Speidel et al. [2019]and splitting into 80% training, 10% validation and 10% test sets. For each scenario we draw mutation rate uniformly: *µ* ∼ *U* (5 × 10^−9^, 2 × 10^−8^) following Smith et al. [2018]; and recombination rate exponentially: *r* ∼ *Exp*(*λ* = 1 × 10^−8^), truncated to [1 × 10^−9^, 1 × 10^−7^]. To account for a large variety of sweep signals, we draw uniformly sweep ages: *τ* ∼ *U* (0, 5, 000) generations; selection strength: *s* ∼ *U* (0.01, 0.05); start frequency saf ∼ *U* (0, 0.1); and end frequency eaf ∼ *U* (0.5, 1).

In addition, similarly to Lauterbur et al. [2023] we tested for both older strong hard sweep and background selection (BGS) to verify previous Flex-sweep results and to set up a proper baseline to test real human data, where the action of BGS and potential sweep candidates due to ancient epidemics is pervasive [Lauterbur et al., 2023, Souilmi et al., 2021, Enard and Petrov, 2020]. We tested both scenarios taking into account YRI-like demography and the same *µ, r, s* and eaf priors.

For the hard old sweep we increased *τ* up to 10,000 human generations ago: *τ* ∼ *U* (5, 000, 10, 000)) accounting for both incomplete and fixed.

To test the presence of BGS, we followed Schrider [2020] central model (see Figure S5). To simplify and speed-up forward-in-time simulations predictions, we first simulated and trained 22,000 neutral and non-neutral (sweeps) regions with discoal following the same prior parameters but under Tennessen et al. [2012] demographic model (Africa 1T129 demographic model provided by stdpopsim). We then use stdpopsim [Adrion et al., 2020] along with the SLiM engine [Haller et al., 2026], to produce 1,000 neutral and non-neutral (BGS) regions following Schrider [2020] central model. We simulated a 50kb non-neutral central region where 75% and 25% were deleterious and neutral mutations respectively. Deleterious mutations were drawn following Kim et al. [2017] values (gamma distribution with mean = −0.013 and shape = 0.186) but decreasing dominance coefficient to *h* = 0.25 accounting for mildly recessive mutations. We avoid rescaling and set-up a burning period of 10. BGS regions were labeled as sweeps during prediction.

### Faster statistics computation

We reimplemented the summary statistics in the feature-vector module taking advantage of BLAS matrix routines, numba just-in-time compilation [Lam et al., 2015] and numpy vectorization. The previous implementation estimated most statistics as explicit loops over pairs of haplotypes, pairs of sites, or focal SNPs, and recomputed several closely intermediate scores independently for each statistic. Main runtime improvements are due to the refactorization of these loops while re-using shared intermediate states across statistics rather than recomputing them. Beyond the explicit numba compilation of individual statistics, we describe below the two main shared optimizations that account for most of the speed-up.

First, several haplotype-based statistics reduce to the same matrix product previously exploited as loops over pairs of haplotypes or pairs of sites. Considering **H** the haplotype matrix of a region, the two products **H**^⊤^**H** and **HH**^⊤^ summarise pairwise allele sharing in the sample. **H** is defined as an *S* × *N* matrix (*S* segregating sites × *N* haplotypes), so that for a site *i* and haplotype *l, H*_*li*_ = 1 if that haplotype carries a derived mutation at such position. Considering that **H** is polarized and phased, the per-site derived count derives as *s*_*i*_ = ∑_*l*_ *H*_*li*_ and the per-haplotype count as *h*_*l*_ = ∑_*i*_ *H*_*li*_. We take advantage of such structure to leverage a single matrix multiplication through BLAS linear-algebra routines to replace the previous pairwise loops, where each off-diagonal entry of **H**^⊤^**H** counts the derived variants that two haplotypes carry in common, while **HH**^⊤^ counts the haplotypes carrying the derived allele at both of a pair of sites.

We used such strategy at three different levels: the haplotype-frequency statistics *HAF* and *iSAFE* [Ronen et al., 2015, Akbari et al., 2018]; the moments of the pairwise haplotype-mismatch distribution following diploS/HIC implementation [Kern and Schrider, 2018] and the *r*^2^ linkage disequilibrium matrix used by *ω*_max_ [Kim and Nielsen, 2004].

The haplotype allele frequency (HAF) score assigned to a haplotype is the sum of the derived-allele counts *s*_*i*_ over the variants it carries [Ronen et al., 2015]; haplotypes that have hitchhiked with a sweep accumulate high-frequency derived alleles and therefore take large HAF values. For haplotype *l* this score is a row sum of the matrix product and a row sum, with monomorphic sites discarded beforehand.

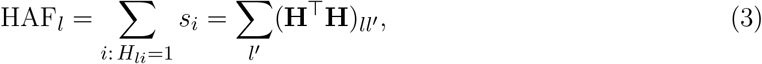

Note, we used the modified HAF of Lauterbur et al. [2023], which reports only the summed upper 10% of the sorted scores.

iSAFE builds on the same HAF quantity to rank the individual variants of a region by how strongly each one carries the sweep signal, separating the candidate favoured variant from neutral hitchhikers [Akbari et al., 2018]. It slides overlapping windows along the region, scores the variants within each window, and combines the windows into a single per-variant ranking. Because every window is scored from the same haplotype matrix product of Eq. 3, we evaluate that product once and reuse it across all windows, compile the per-window scoring with numba, and perform the grouping and ranking of variants across windows as a single polars aggregation.

Following diploS/HIC [Kern and Schrider, 2018], we summarise the distribution of pairwise mismatch distances *d*_*ij*_ (the number of sites at which haplotypes *i* and *j* differ) by its variance, skewness and kurtosis. Again we used the **H**^*T*^ **H** shared product to capture the distance and the mismatch matrix to get the three moments in one single vectorized operation

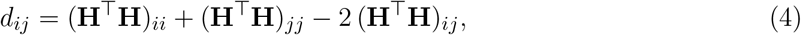

*ω*_max_ detects a sweep through the excess of linkage disequilibrium it builds between the two flanks of the selected site [Kim and Nielsen, 2004]. We computed from the *r*^2^ matrix values following the same site-by-site product **HH**^⊤^ over genotypes and compute *ω*_max_ through numba optimzation by maximizing the ratio of within-to between-flank linkage disequilibrium for each focal SNP.

Second, we reimplemented *DIND* [Barreiro et al., 2009] and the original Flex-sweep statistics (*Sratio, highfreq, lowfreq, hapDAF* -*o* and *hapDAF* -*s*) [Lauterbur et al., 2023], which quantify how a sweep distorts the variation flanking a focal SNP by contrasting its derived against its ancestral background. A single numba kernel identifies the focal SNPs of intermediate frequency (0.25 ≤ *f* ≤ 0.95), locates each flanking window by binary search over the sorted positions, and splits the haplotypes into the two backgrounds while accumulating, for every neighbouring variant, its frequency on the derived background (*f*_*d*_), on the ancestral background (*f*_*a*_), and its overall frequency (*f*_*tot*_). Pre-computed {*f*_*d*_, *f*_*a*_, *f*_*tot*_} value are then reused as needed to derive all six statistics at once through vectorized numpy reductions.

### Domain-adaptive gradient schedule

Rather than applying a fixed gradient-reversal factor, we stabilize the adversarial adaptation through controlled gradient scheduling, so that the model first learns task-discriminative features from the labeled simulations before enforcing domain invariance. This prevents the domain discriminator from distorting the feature extractor early in training, when simulation-specific artifacts would otherwise dominate the gradient. Accordingly, the gradient-reversal strength follows an epoch-dependent ramp from zero to a predefined maximum,

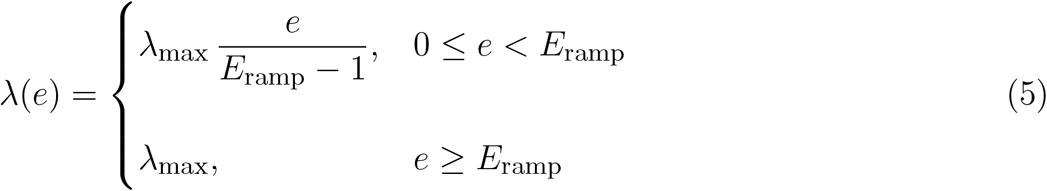

where *e* is the epoch, *E*_ramp_ the number of warm-up epochs, and *λ*_max_ the target GRL strength. We cap the adversarial strength at *λ*_max_ to keep the domain objective from dominating optimization, reducing negative transfer and preserving features that generalize across mismatched empirical and simulated distributions.

**Fig. S1:**
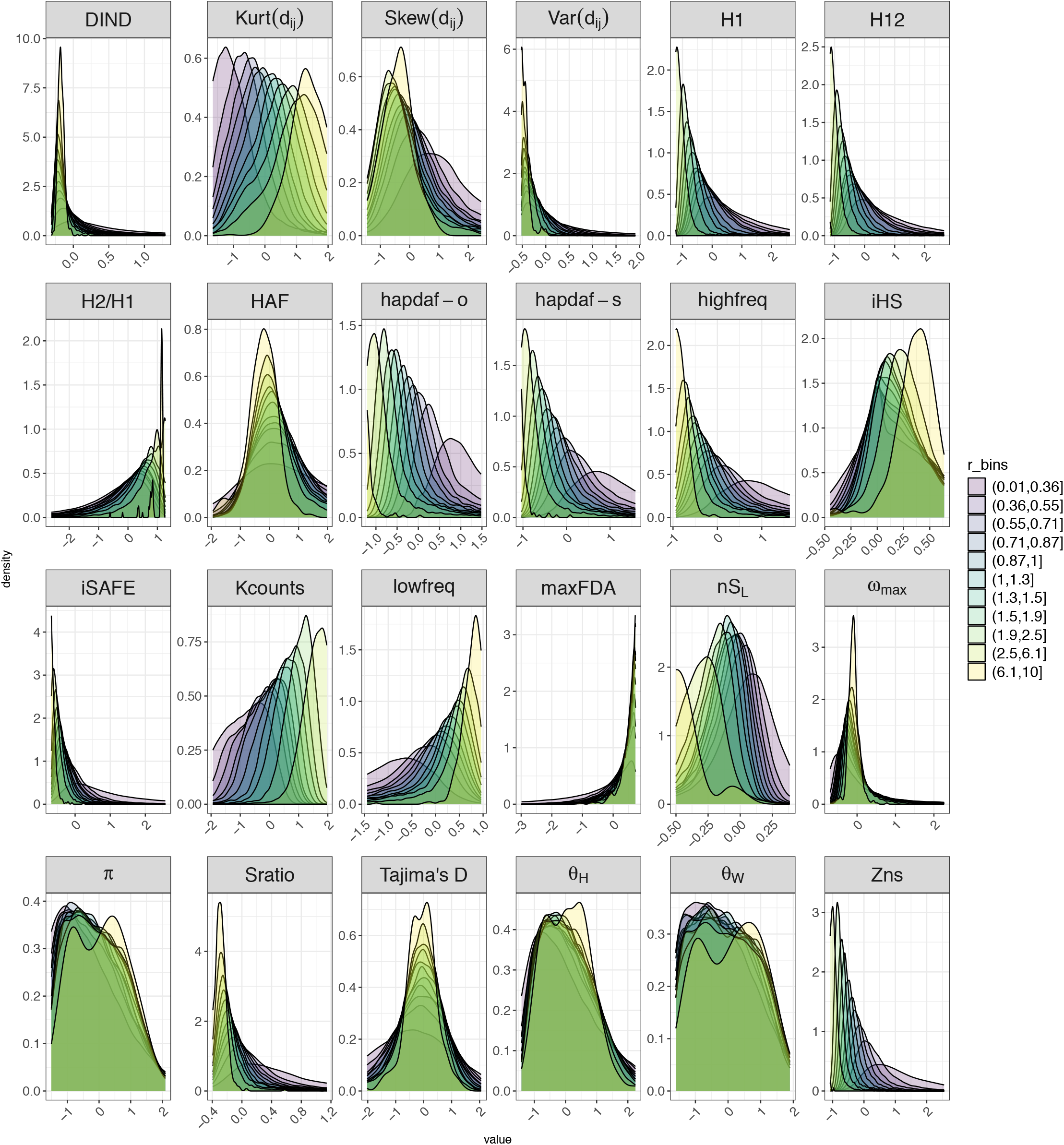
Flex-sweep Z-scored summary statistics distribution by recombination bin simulated

**Fig. S2:**
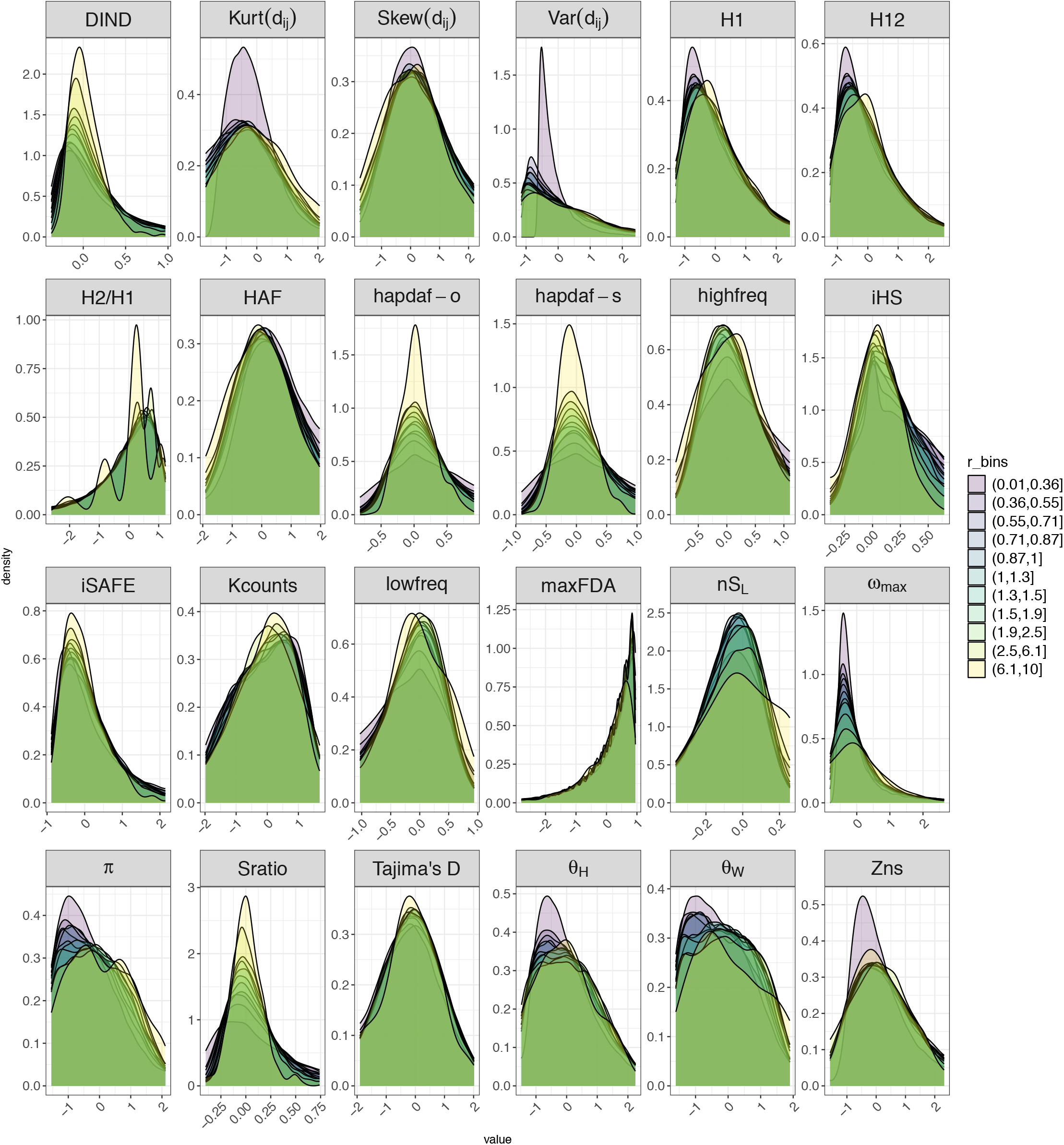
Flex-sweep recombination stratified Z-scored summary statistics distribution by recombination bin simulated

**Fig. S3:**
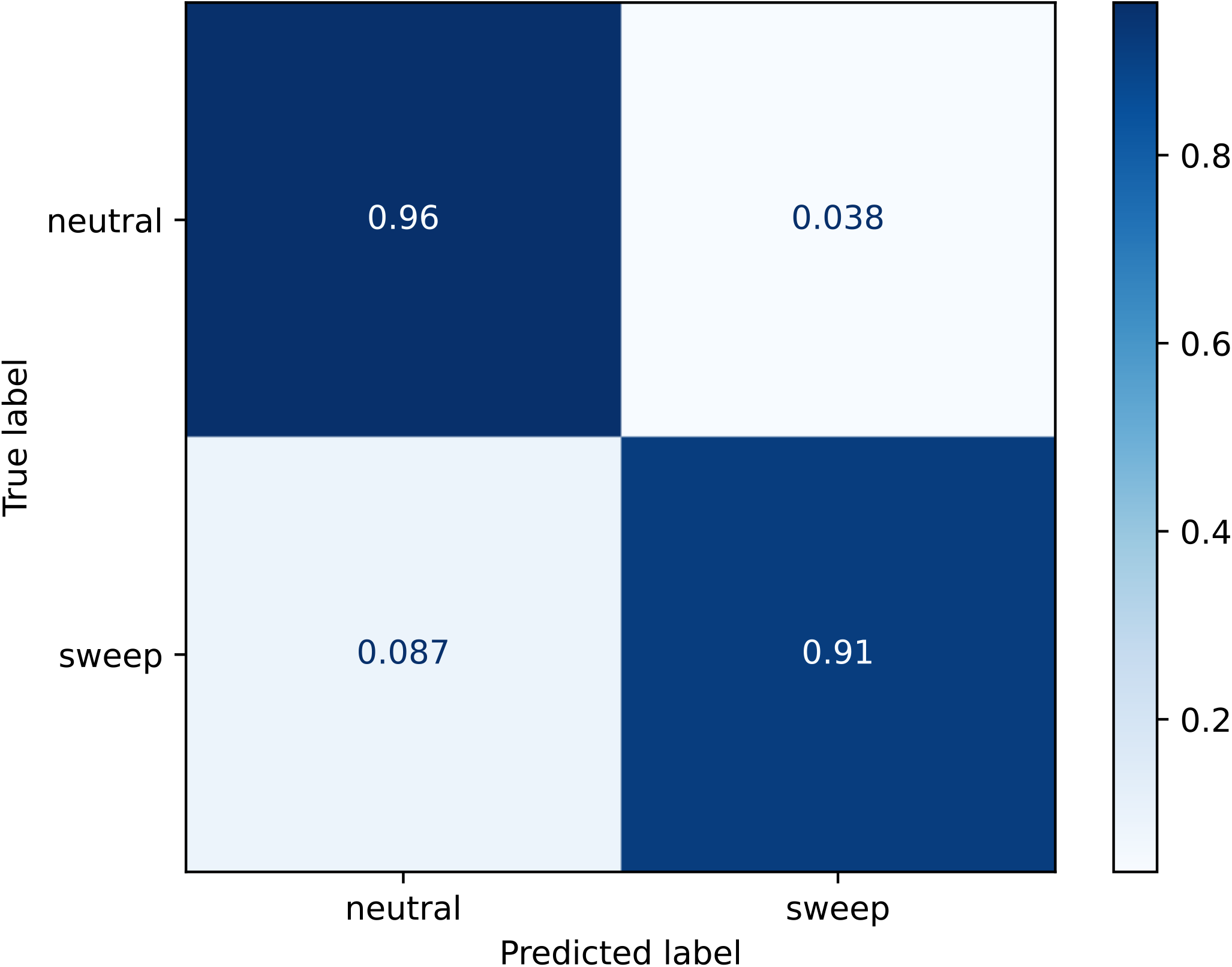
YRI confusion matrix on 250,000 neutral and non-neutral trained dataset.

**Fig. S4:**
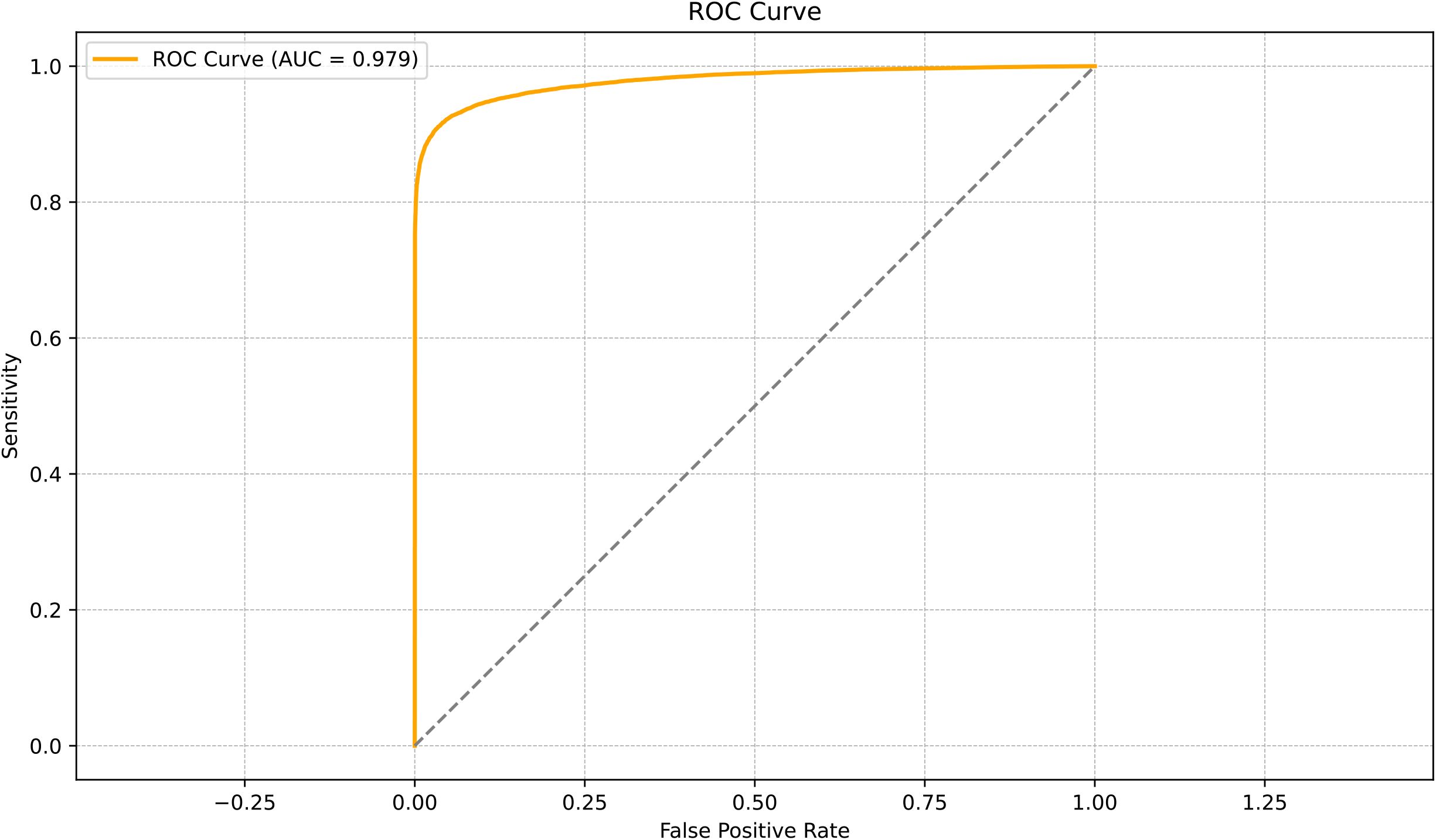
YRI ROC curve on 250,000 neutral and non-neutral trained dataset.

**Fig. S5:**
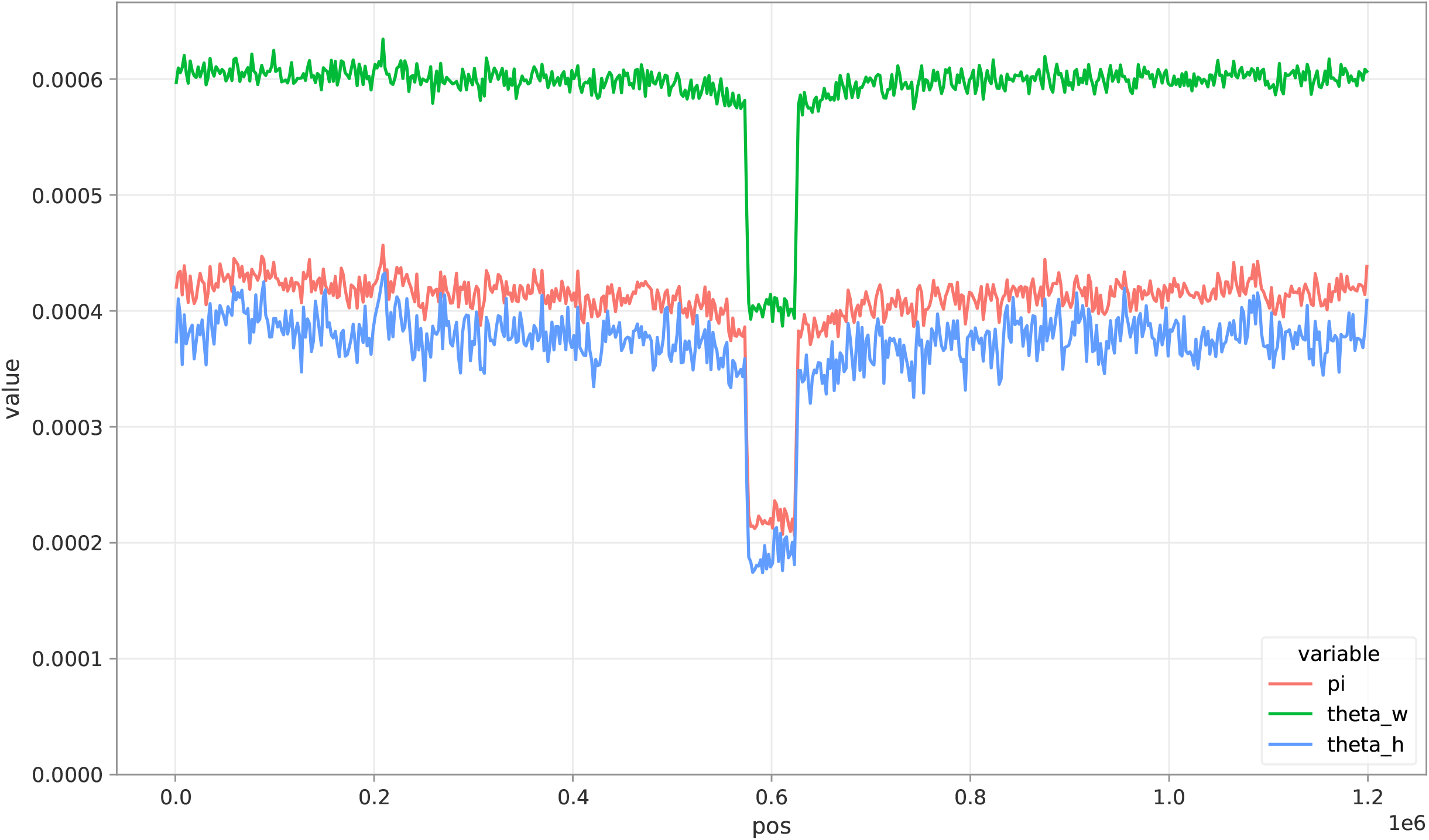
Central Background selection model from Schrider [2020]. 50Kb middle region following Kim et al. [2017] DFE with midly recessive dominance (*h* = 0.25).

**Fig. S6:**
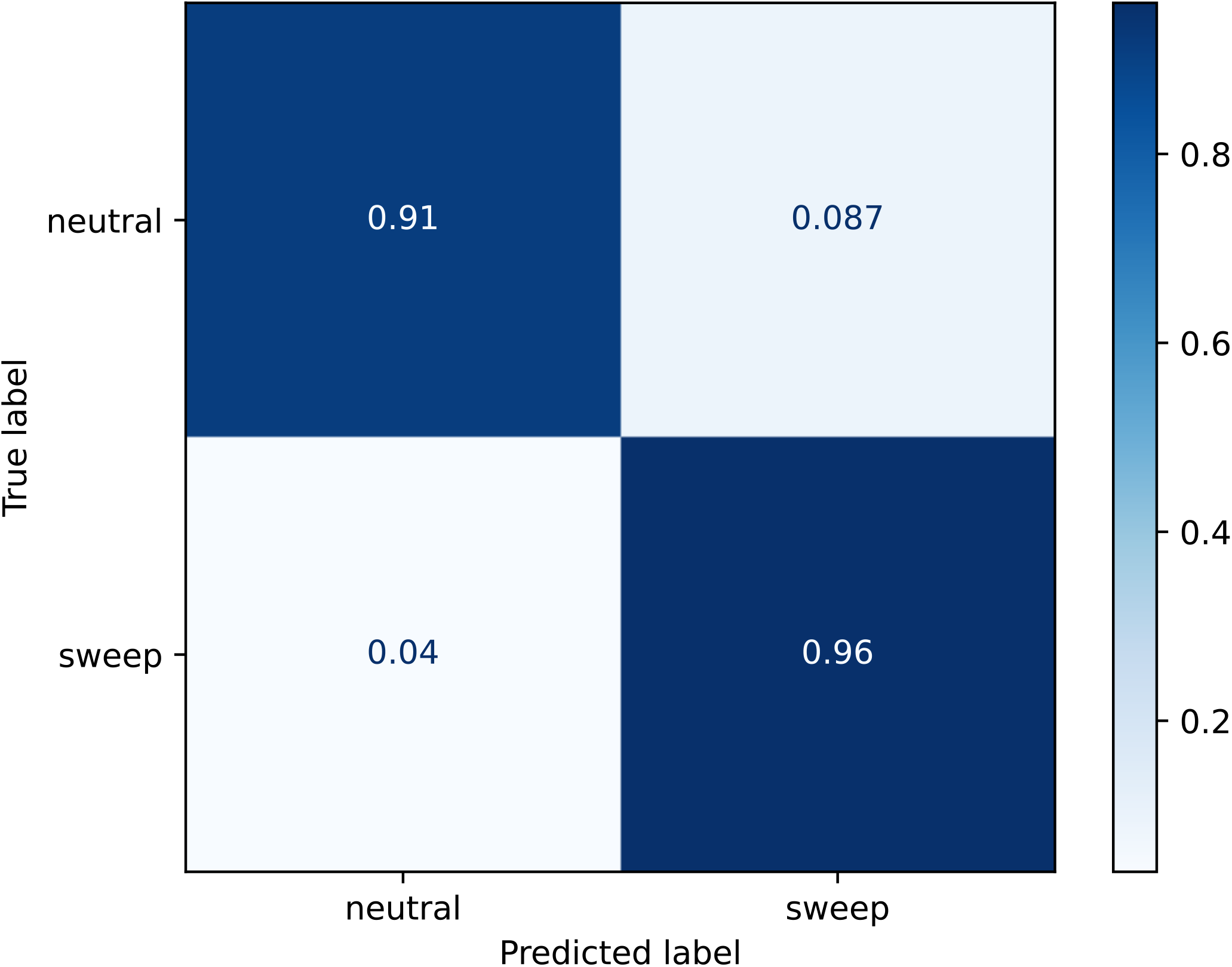
Confusion matrix for hard old sweep set. 1,000 neutral and non-neutral simulations following YRI-like demography were tested accounting for timing distribution *t U* (5, 000, 10, 000) while increasing end allele frequency to [0.75, 1]. The prediction was performed using the baseline trained CNN accounting for 250,000 neutral and non-neutral regions with timing distribution of *t U* (0, 5000).

**Fig. S7:**
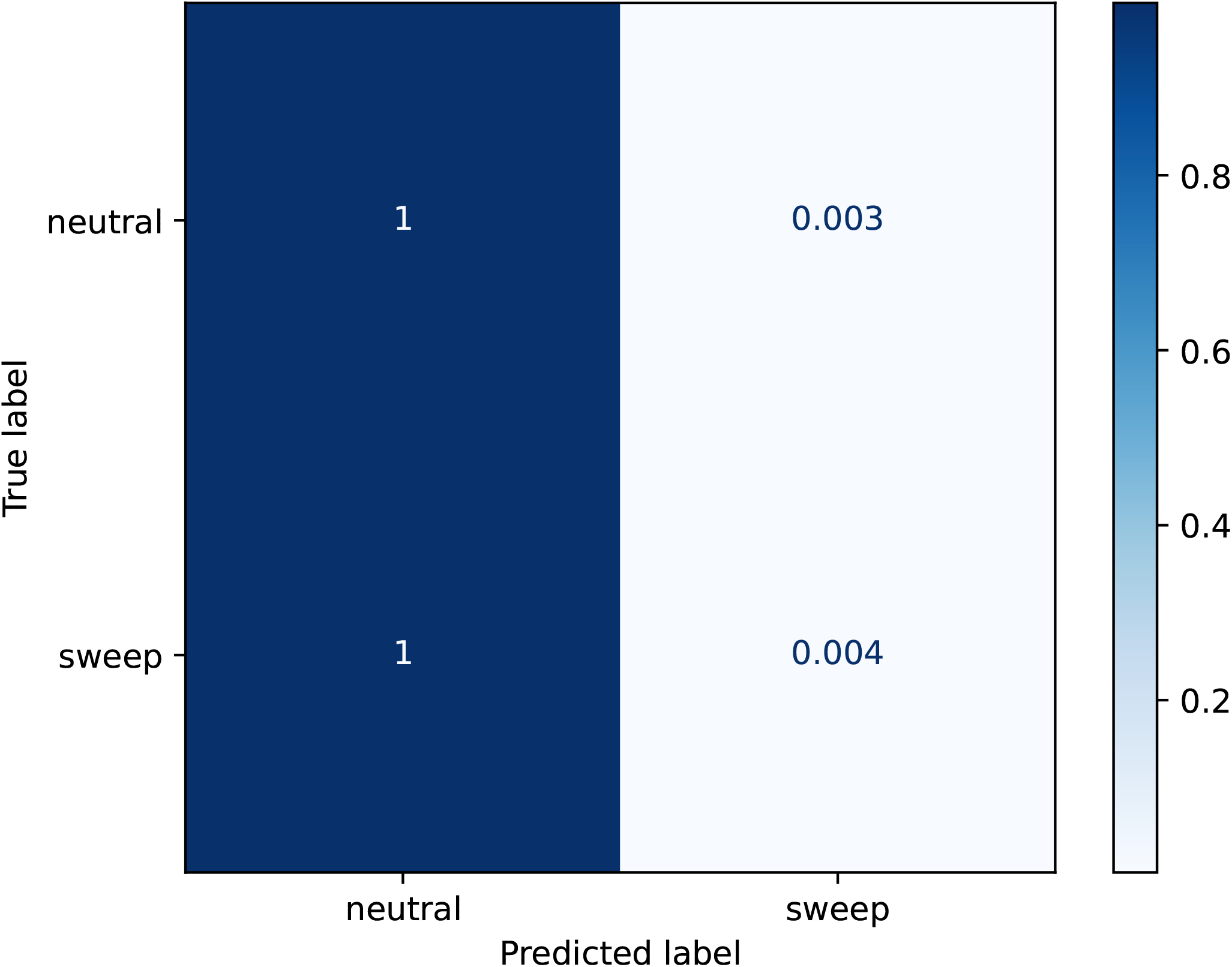
Confusion matrix for background selection (BGS) simulations. 1,000 neutral and non-neutral simulations were performed using stdpopsim under the Tennesen model. BGS were simulated under the Schrider and Kern [2017] central model following Kim et al. [2017] DFE. BGS simulations were labelled as sweeps.

**Fig. S8:**
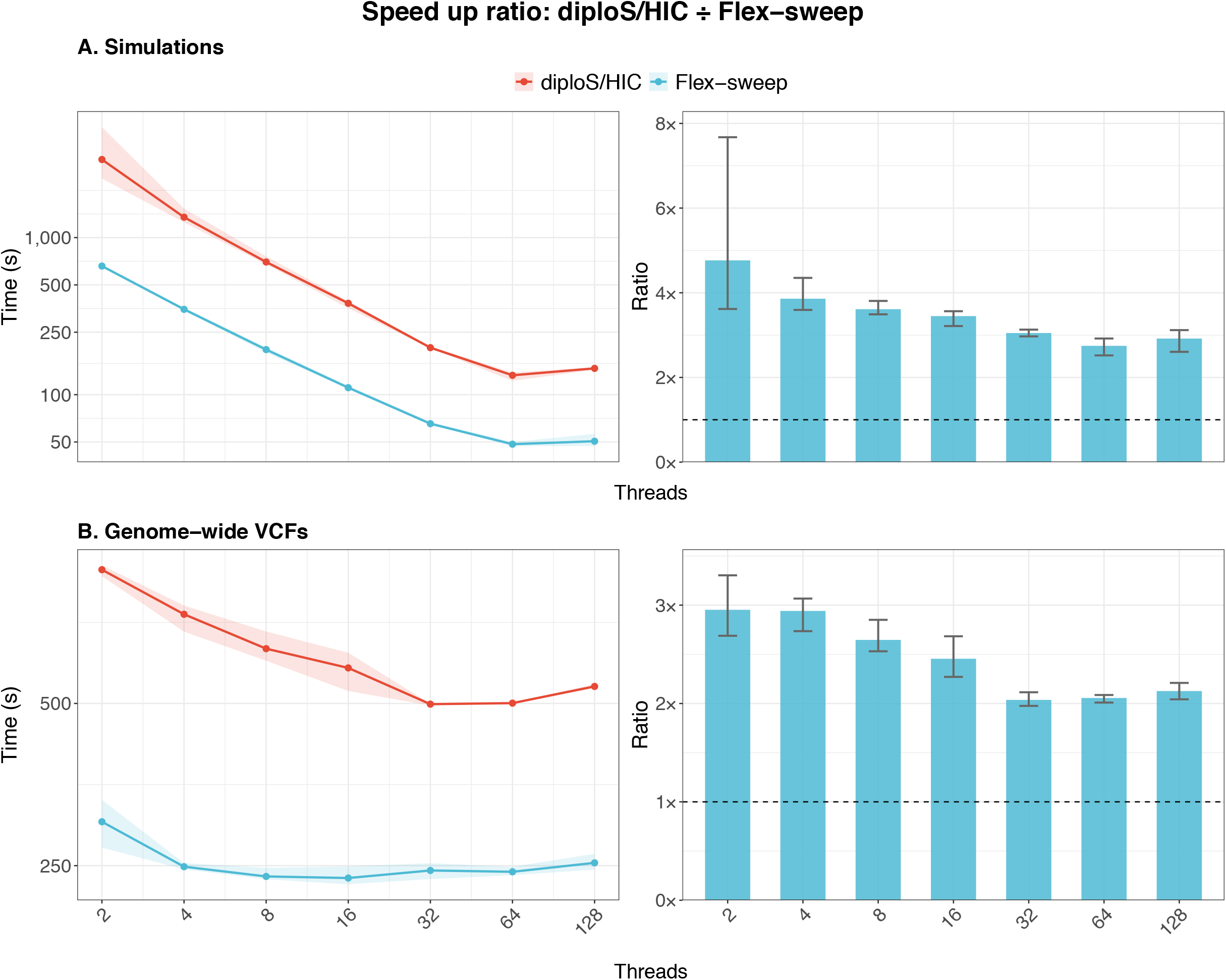
Flex-sweep benchmark. A. Runtime comparison on 22,000 simulations neutral and non-neutral simulations. B. Runtime comparison on 1000GP YRI 22 autosomes.

## References

Yuseob Kim and Rasmus Nielsen. Linkage Disequilibrium as a Signature of Selective Sweeps. Genetics, 167(3):1513–1524, July 2004. ISSN 1943-2631. doi: 10.1534/genetics.103.025387. URL https://doi.org/10.1534/genetics.103.025387.

Benjamin F. Voight, Sridhar Kudaravalli, Xiaoquan Wen, and Jonathan K. Pritchard. A Map of Recent Positive Selection in the Human Genome. PLOS Biology, 4(3): e72, March 2006. ISSN 1545-7885. doi: 10.1371/journal.pbio.0040072. URL https://journals.plos.org/plosbiology/article?id=10.1371/journal.pbio.0040072. Publisher: Public Library of Science.

Nandita R. Garud, Philipp W. Messer, Erkan O. Buzbas, and Dmitri A. Petrov. Recent Selective Sweeps in North American Drosophila melanogaster Show Signatures of Soft Sweeps. PLOS Genetics, 11(2):e1005004, February 2015. ISSN 1553-7404. doi: 10.1371/journal.pgen.1005004. URL https://journals.plos.org/plosgenetics/article?id=10.1371/journal.pgen.1005004. Publisher: Public Library of Science.

Ali Akbari, Joseph J. Vitti, Arya Iranmehr, Mehrdad Bakhtiari, Pardis C. Sabeti, Siavash Mirarab, and Vineet Bafna. Identifying the favored mutation in a positive selective sweep. Nature Methods, 15(4):279–282, April 2018. ISSN 1548-7105. doi: 10.1038/nmeth.4606. URL https://www.nature.com/articles/nmeth.4606. Publisher: Nature Publishing Group.

Scott H. Williamson, Melissa J. Hubisz, Andrew G. Clark, Bret A. Payseur, Carlos D. Bustamante, and Rasmus Nielsen. Localizing Recent Adaptive Evolution in the Human Genome. PLOS Genetics, 3(6):e90, June 2007. ISSN 1553-7404. doi: 10.1371/journal.pgen.0030090. URL https://journals.plos.org/plosgenetics/article?id=10.1371/journal.pgen.0030090. Publisher: Public Library of Science.

Michael DeGiorgio, Christian D. Huber, Melissa J. Hubisz, Ines Hellmann, and Rasmus Nielsen. SweepFinder2: increased sensitivity, robustness and flexibility. Bioinformatics, 32 (12):1895–1897, June 2016. ISSN 1367-4803. doi: 10.1093/bioinformatics/btw051. URL https://doi.org/10.1093/bioinformatics/btw051.

Alexandre M Harris and Michael DeGiorgio. A Likelihood Approach for Uncovering Selective Sweep Signatures from Haplotype Data. Molecular Biology and Evolution, 37(10): 3023–3046, October 2020. ISSN 0737-4038. doi: 10.1093/molbev/msaa115. URL https://doi.org/10.1093/molbev/msaa115.

Sharon R. Grossman, Ilya Shylakhter, Elinor K. Karlsson, Elizabeth H. Byrne, Shannon Morales, Gabriel Frieden, Elizabeth Hostetter, Elaine Angelino, Manuel Garber, Or Zuk, Eric S. Lander, Stephen F. Schaffner, and Pardis C. Sabeti. A Composite of Multiple Signals Distinguishes Causal Variants in Regions of Positive Selection. Science, 327(5967):883–886, February 2010. doi: 10.1126/science.1183863. URL https://www.science.org/doi/10.1126/science.1183863. Publisher: American Association for the Advancement of Science.

Lauren Alpert Sugden, Elizabeth G. Atkinson, Annie P. Fischer, Stephen Rong, Brenna M. Henn, and Sohini Ramachandran. Localization of adaptive variants in human genomes using averaged one-dependence estimation. Nature Communications, 9(1):703, February 2018. ISSN 2041-1723. doi: 10.1038/s41467-018-03100-7. URL https://www.nature.com/articles/s41467-018-03100-7. Publisher: Nature Publishing Group.

Nikolaos Alachiotis and Pavlos Pavlidis. RAiSD detects positive selection based on multiple signatures of a selective sweep and SNP vectors. Communications Biology, 1(1):79, June 2018. ISSN 2399-3642. doi: 10.1038/s42003-018-0085-8. URL https://www.nature.com/articles/s42003-018-0085-8. Publisher: Nature Publishing Group.

Benjamin M. Peter, Emilia Huerta-Sanchez, and Rasmus Nielsen. Distinguishing between Selective Sweeps from Standing Variation and from a De Novo Mutation. PLOS Genetics, 8(10):e1003011, October 2012. ISSN 1553-7404. doi: 10.1371/journal.pgen.1003011. URL https://journals.plos.org/plosgenetics/article?id=10.1371/journal.pgen.1003011. Publisher: Public Library of Science.

Fernando Racimo, Martin Kuhlwilm, and Montgomery Slatkin. A Test for Ancient Selective Sweeps and an Application to Candidate Sites in Modern Humans. Molecular Biology and Evolution, 31(12):3344–3358, December 2014. ISSN 0737-4038. doi: 10.1093/molbev/msu255. URL https://doi.org/10.1093/molbev/msu255.

Parul Johri, Susanne P Pfeifer, and Jeffrey D Jensen. Developing an Evolutionary Baseline Model for Humans: Jointly Inferring Purifying Selection with Population History. Molecular Biology and Evolution, 40(5):msad100, May 2023. ISSN 1537-1719. doi: 10.1093/molbev/msad100. URL https://doi.org/10.1093/molbev/msad100.

Pavlos Pavlidis, Jeffrey D Jensen, and Wolfgang Stephan. Searching for Footprints of Positive Selection in Whole-Genome SNP Data From Nonequilibrium Populations. Genetics, 185(3): 907–922, July 2010. ISSN 1943-2631. doi: 10.1534/genetics.110.116459. URL https://doi.org/10.1534/genetics.110.116459.

Andrew D Kern and Daniel R Schrider. diploS/HIC: An Updated Approach to Classifying Selective Sweeps. G3 Genes|Genomes|Genetics, 8(6):1959–1970, June 2018. ISSN 2160-1836. doi: 10.1534/g3.118.200262. URL https://doi.org/10.1534/g3.118.200262.

Hussein A Hejase, Ziyi Mo, Leonardo Campagna, and Adam Siepel. A Deep-Learning Approach for Inference of Selective Sweeps from the Ancestral Recombination Graph. Molecular Biology and Evolution, 39(1):msab332, January 2022. ISSN 1537-1719. doi: 10.1093/molbev/msab332. URL https://doi.org/10.1093/molbev/msab332.

Ziyi Mo and Adam Siepel. Domain-adaptive neural networks improve supervised machine learning based on simulated population genetic data. PLOS Genetics, 19(11):e1011032, November 2023. ISSN 1553-7404. doi: 10.1371/journal.pgen.1011032. URL https://journals.plos.org/plosgenetics/article?id=10.1371/journal.pgen.1011032. Publisher: Public Library of Science.

Sjoerd van den Belt and Nikolaos Alachiotis. Fast and accurate deep learning scans for signatures of natural selection in genomes using FASTER-NN. Communications Biology, 8 (1):58, January 2025. ISSN 2399-3642. doi: 10.1038/s42003-025-07480-7. URL https://www.nature.com/articles/s42003-025-07480-7. Publisher: Nature Publishing Group.

Logan S Whitehouse, Dylan D Ray, and Daniel R Schrider. Tree Sequences as a General-Purpose Tool for Population Genetic Inference. Molecular Biology and Evolution, 41(11):msae223, November 2024. ISSN 1537-1719. doi: 10.1093/molbev/msae223. URL https://doi.org/10.1093/molbev/msae223.

Sandipan Paul Arnab, Andre Luiz Campelo dos Santos, Matteo Fumagalli, and Michael DeGiorgio. Efficient Detection and Characterization of Targets of Natural Selection Using Transfer Learning. Molecular Biology and Evolution, 42(5):msaf094, May 2025. ISSN 1537-1719. doi: 10.1093/molbev/msaf094. URL https://doi.org/10.1093/molbev/msaf094.

Hanqing Zhao and Nikolaos Alachiotis. Data preprocessing methods for selective sweep detection using convolutional neural networks. Methods, 233:19–29, January 2025. ISSN 1046-2023. doi: 10.1016/j.ymeth.2024.11.003. URL https://www.sciencedirect.com/science/article/pii/S1046202324002408.

Ian V. Caldas, Andrew G. Clark, and Philipp W. Messer. Inference of selective sweep parameters through supervised learning, July 2022. URL https://www.biorxiv.org/content/10.1101/2022.07.19.500702v1. Pages: 2022.07.19.500702 Section: New Results.

M Elise Lauterbur, Kasper Munch, and David Enard. Versatile Detection of Diverse Selective Sweeps with Flex-Sweep. Molecular Biology and Evolution, 40(6):msad139, June 2023. ISSN 1537-1719. doi: 10.1093/molbev/msad139. URL https://doi.org/10.1093/molbev/msad139.

Felix M. Key, Qiaomei Fu, Frédéric Romagné, Michael Lachmann, and Aida M. Andrés. Human adaptation and population differentiation in the light of ancient genomes. Nature Communications, 7(1):10775, March 2016. ISSN 2041-1723. doi: 10.1038/ncomms10775. URL https://www.nature.com/articles/ncomms10775. Publisher: Nature Publishing Group.

Fernando Racimo. Testing for Ancient Selection Using Cross-population Allele Frequency Differentiation. Genetics, 202(2): 733–750, February 2016. ISSN 1943-2631. doi: 10.1534/genetics.115.178095. URL https://doi.org/10.1534/genetics.115.178095.

Xiaoheng Cheng, Cheng Xu, and Michael DeGiorgio. Fast and robust detection of ancestral selective sweeps. Molecular Ecology, 26(24):6871–6891, 2017. ISSN 1365-294X. doi: 10.1111/mec.14416. URL https://onlinelibrary.wiley.com/doi/abs/10.1111/mec.14416. eprint: https://onlinelibrary.wiley.com/doi/pdf/10.1111/mec.14416.

Stéphane Peyrégne, Michael James Boyle, Michael Dannemann, and Kay Prüfer. Detecting ancient positive selection in humans using extended lineage sorting. Genome Research, 27(9):1563– 1572, September 2017. ISSN 1088-9051, 1549-5469. doi: 10.1101/gr.219493.116. URL http://genome.cshlp.org/content/27/9/1563. Company: Cold Spring Harbor Laboratory Press Distributor: Cold Spring Harbor Laboratory Press Institution: Cold Spring Harbor Laboratory Press Label: Cold Spring Harbor Laboratory Press Publisher: Cold Spring Harbor Lab.

Ryan D. Hernandez, Joanna L. Kelley, Eyal Elyashiv, S. Cord Melton, Adam Auton, Gilean McVean, 1000 GENOMES PROJECT, Guy Sella, and Molly Przeworski. Classic Selective Sweeps Were Rare in Recent Human Evolution. Science, 331(6019):920–924, February 2011. doi: 10.1126/science.1198878. URL https://www.science.org/doi/10.1126/science.1198878.

Marta Byrska-Bishop, Uday S. Evani, Xuefang Zhao, Anna O. Basile, Haley J. Abel, Allison A. Regier, André Corvelo Wayne E. Clarke, Rajeeva Musunuri, Kshithija Nagulapalli, Susan Fairley, Alexi Runnels, Lara Winterkorn, Ernesto Lowy, Evan E. Eichler, Jan O. Korbel, Charles Lee, Tobias Marschall, Scott E. Devine, William T. Harvey, Weichen Zhou, Ryan E. Mills, Tobias Rausch, Sushant Kumar, Can Alkan, Fereydoun Hormozdiari, Zechen Chong, Yu Chen, Xiaofei Yang, Jiadong Lin, Mark B. Gerstein, Ye Kai, Qihui Zhu, Feyza Yilmaz, Chunlin Xiao, Paul Flicek, Soren Germer, Harrison Brand, Ira M. Hall, Michael E. Talkowski, Giuseppe Narzisi, and Michael C. Zody.High-coverage whole-genome sequencing of the expanded 1000 Genomes Project cohort including 602 trios. Cell, 185(18):3426–3440.e19, September 2022. ISSN 0092-8674, 1097-4172. doi: 10.1016/j.cell.2022.08.004. URL https://www.cell.com/cell/abstract/S0092-8674(22)00991-6. Publisher: Elsevier.

Graham Gower, Aaron P Ragsdale, Gertjan Bisschop, Ryan N Gutenkunst, Matthew Hartfield, Ekaterina Noskova, Stephan Schiffels, Travis J Struck, Jerome Kelleher, and Kevin R Thornton. Demes: a standard format for demographic models. Genetics, 222(3):iyac131, November 2022. ISSN 1943-2631. doi: 10.1093/genetics/iyac131. URL https://doi.org/10.1093/genetics/iyac131.

Andrew D. Kern and Daniel R. Schrider. coalescent simulations with selection. Bioinformatics, 32 (24):3839–3841, December 2016. ISSN 1367-4803. doi: 101093/bioinformatics/btw556. URL https://doi.org/10.1093/bioinformatics/btw556.

Siu Kwan Lam, Antoine Pitrou, and Stanley Seibert. Numba: a LLVM-based Python JIT compiler. In Proceedings of the Second Workshop on the LLVM Compiler Infrastructure in HPC, pages 1–6, Austin Texas, November 2015. ACM. ISBN 978-1-4503-4005-2. doi: 10.1145/2833157.2833162. URL https://dl.acm.org/doi/10.1145/2833157.2833162.

Luis B. Barreiro, Meriem Ben-Ali, Hélène Quach, Guillaume Laval, Etienne Patin, Joseph K. Pickrell, Christiane Bouchier, Magali Tichit, Olivier Neyrolles, Brigitte Gicquel, Judith R. Kidd, Kenneth K. Kidd, Alexandre Alcaïs, Josiane Ragimbeau, Sandra Pellegrini, Laurent Abel, Jean-Laurent Casanova, and Llúis Quintana-Murci. Evolutionary Dynamics of Human Toll-Like Receptors and Their Different Contributions to Host Defense. PLOS Genetics, 5(7):e1000562, July 2009. ISSN 1553-7404. doi: 10.1371/journal.pgen.1000562. URL https://journals.plos.org/plosgenetics/article?id=10.1371/journal.pgen.1000562. Publisher: Public Library of Science.

Roy Ronen, Glenn Tesler, Ali Akbari, Shay Zakov, Noah A. Rosenberg, and Vineet Bafna. Predicting Carriers of Ongoing Selective Sweeps without Knowledge of the Favored Allele. PLOS Genetics, 11(9):e1005527, September 2015. ISSN 1553-7404. doi: 10.1371/journal.pgen.1005527. URL https://journals.plos.org/plosgenetics/article?id=10.1371/journal.pgen.1005527. Publisher: Public Library of Science.

Ritchie Vink, Alexander Beedienameexhaustion, Orson Peters, Gijs Burghoorn, Stijn de Gooijer, Marco Edward Gorelli, reswqa, Marshall, Jeroen van Zundert, Gert Hulselmans, Koen Denecker, Cory Grinstead, Luke Manley, Amber SprenkelschielP, Kevin Patyk, Itamar Turner-Trauring, Kuba Valtar, Lawrence Mitchell, Karl Genockey, Henry Harbeck eitsupi, Lukas Bergdoll, Robin, deanm 0000, Oliver Borchert, Matteo Santamaria, and Ion Koutsouris. pola-rs/polars: Python Polars 1.39.3, March 2026. URL https://zenodo.org/records/19130590.

Joshua M. Akey. Constructing genomic maps of positive selection in humans: Where do we go from here? Genome Research, 19 (5):711–722, May 2009. ISSN 1088-9051, 1549-5469. doi: 10.1101/gr.086652.108. URL http://genome.cshlp.org/content/19/5/711.

Ulas Isildak, Alessandro Stella, and Matteo Fumagalli. Distinguishing between recent balancing selection and incomplete sweep using deep neural networks. Molecular Ecology Resources, 21(8):2706–2718, 2021. ISSN 1755-0998. doi: 10.1111/1755-0998.13379. URL https://onlinelibrary.wiley.com/doi/abs/10.1111/1755-0998.13379. eprint: https://onlinelibrary.wiley.com/doi/pdf/10.1111/1755-0998.13379.

Tom R. Booker, Sam Yeaman, and Michael C. Whitlock. Variation in recombination rate affects detection of outliers in genome scans under neutrality. Molecular Ecology, 29(22):4274–4279, 2020. ISSN 1365-294X. doi: 10.1111/mec.15501. URL https://onlinelibrary.wiley.com/doi/abs/10.1111/mec.15501. eprint: https://onlinelibrary.wiley.com/doi/pdf/10.1111/mec.15501.

Kelsey Elizabeth Johnson and Benjamin F. Voight. Patterns of shared signatures of recent positive selection across human populations. Nature Ecology & Evolution, 2(4):713–720, April 2018. ISSN 2397-334X. doi: 10.1038/s41559-018-0478-6. URL https://www.nature.com/articles/s41559-018-0478-6.

Jun Ishigohoka and Miriam Liedvogel. High-recombining genomic regions affect demography inference based on ancestral recombination graphs. Genetics, 229(3):iyaf004, March 2025. ISSN 1943-2631. doi: 10.1093/genetics/iyaf004. URL https://doi.org/10.1093/genetics/iyaf004.

Bjarni V. Halldorsson, Gunnar Palsson, Olafur A. Stefansson, Hakon Jonsson, Marteinn T. Hardarson, Hannes P. Eggertsson, Bjarni Gunnarsson, Asmundur Oddsson, Gisli H. Halldorsson, Florian Zink, Sigurjon A. Gudjonsson, Michael L. Frigge, Gudmar Thorleifsson, Asgeir Sigurdsson, Simon N. Stacey, Patrick Sulem, Gisli Masson, Agnar Helgason, Daniel F. Gudbjartsson, Unnur Thorsteinsdottir, and Kari Stefansson. Characterizing mutagenic effects of recombination through a sequence-level genetic map. Science, 363(6425):eaau1043, January 2019. doi: 10.1126/science.aau1043. URL https://www.science.org/doi/abs/10.1126/science.aau1043.

Linh N Tran, David Castellano, and Ryan N Gutenkunst. Interpreting Supervised Machine Learning Inferences in Population Genomics Using Haplotype Matrix Permutations. Molecular Biology and Evolution, 42(10):msaf250, October 2025. ISSN 1537-1719. doi: 10.1093/molbev/msaf250. URL https://doi.org/10.1093/molbev/msaf250.

Ryan D. Hernandez, Scott H. Williamson, and Carlos D. Bustamante. Context Dependence, Ancestral Misidentification, and Spurious Signatures of Natural Selection. Molecular Biology and Evolution, 24(8):1792–1800, August 2007. ISSN 0737-4038. doi: 10.1093/molbev/msm108. URL https://doi.org/10.1093/molbev/msm108.

Peter D Keightley and Benjamin C Jackson. Inferring the Probability of the Derived vs. the Ancestral Allelic State at a Polymorphic Site. Genetics, 209(3):897–906, July 2018. ISSN 1943-2631. doi: 10.1534/genetics.118.301120. URL https://doi.org/10.1534/genetics.118.301120.

Vince Buffalo. vsbuffalo/maftk, December 2025. URL https://github.com/vsbuffalo/maftk.original-date:2024-11-22T07:20:35Z.

Joel Armstrong, Glenn Hickey, Mark Diekhans, Ian T. Fiddes, Adam M. Novak, Alden Deran, Qi Fang, Duo Xie, Shaohong Feng, Josefin Stiller, Diane Genereux, Jeremy Johnson, Voichita Dana Marinescu, Jessica Alföldi, Robert S. Harris, Kerstin Lindblad-Toh, David Haussler, Elinor Karlsson, Erich D. Jarvis, Guojie Zhang, and Benedict Paten. Progressive Cactus is a multiple-genome aligner for the thousand-genome era. Nature, 587(7833):246–251, November 2020. ISSN 1476-4687. doi: 10.1038/s41586-020-2871-y. URL https://www.nature.com/articles/s41586-020-2871-y.

Lukas F. K. Kuderna, Jacob C. Ulirsch, Sabrina Rashid, Mohamed Ameen, Laksshman Sundaram, Glenn Hickey, Anthony J. Cox, Hong Gao, Arvind Kumar, Francois Aguet, Matthew J. Christmas, Hiram Clawson, Maximilian Haeussler, Mareike C. Janiak, Martin Kuhlwilm, Joseph D. Orkin, Thomas Bataillon, Shivakumara Manu, Alejandro Valenzuela, Juraj Bergman, Marjolaine Rouselle, Felipe Ennes Silva, Lidia Agueda, Julie Blanc, Marta Gut, Dorien de Vries, Ian Goodhead, R. Alan Harris, Muthuswamy Raveendran, Axel Jensen, Idriss S. Chuma, Julie E. Horvath, Christina Hvilsom, David Juan, Peter Frandsen, Joshua G. Schraiber, Fabiano R. de Melo, Fabrício Bertuol, Hazel Byrne, Iracilda Sampaio, Izeni Farias, João Valsecchi, Malu Messias, Maria N. F. da Silva, Mihir Trivedi, Rogerio Rossi, Tomas Hrbek, Nicole Andriaholinirina, Clément J. Rabarivola, Alphonse Zaramody, Clifford J. Jolly, Jane Phillips-Conroy, Gregory Wilkerson, Christian Abee, Joe H. Simmons, Eduardo Fernandez-Duque, Sree Kanthaswamy, Fekadu Shiferaw, Dongdong Wu, Long Zhou, Yong Shao, Guojie Zhang, Julius D. Keyyu, Sascha Knauf, Minh D. Le, Esther Lizano, Stefan Merker, Arcadi Navarro, Tilo Nadler, Chiea Chuen Khor, Jessica Lee, Patrick Tan, Weng Khong Lim, Andrew C. Kitchener, Dietmar Zinner, Ivo Gut, Amanda D. Melin, Katerina Guschanski, Mikkel Heide Schierup, Robin M. D. Beck, Ioannis Karakikes, Kevin C. Wang, Govindhaswamy Umapathy, Christian Roos, Jean P. Boubli, Adam Siepel, Anshul Kundaje, Benedict Paten, Kerstin Lindblad-Toh, Jeffrey Rogers, Tomas Marques Bonet, and Kyle Kai-How Farh. Identification of constrained sequence elements across 239 primate genomes. Nature, 625(7996): 735–742, January 2024. ISSN 1476-4687. doi: 10.1038/s41586-023-06798-8. URL https://www.nature.com/articles/s41586-023-06798-8.

Marek Wiewiórka, Pavel Khamutou, Marek Zbysiński, and Tomasz Gambin. polars-bio—fast, scalable, and out-of-core operations on large genomic interval datasets. Bioinformatics, 41(12):btaf640, December 2025. ISSN 1367-4811. doi: 10.1093/bioinformatics/btaf640. URL https://doi.org/10.1093/bioinformatics/btaf640.

David Enard and Dmitri A. Petrov. Ancient RNA virus epidemics through the lens of recent adaptation in human genomes. Philosophical Transactions of the Royal Society B: Biological Sciences, 375(1812):20190575, October 2020. doi: 10.1098/rstb.2019.0575. URL https://royalsocietypublishing.org/doi/full/10.1098/rstb.2019.0575. Publisher: Royal Society.

Chenlu Di, Jesus Murga Moreno, Diego F Salazar-Tortosa, M Elise Lauterbur, and David Enard. Decreased recent adaptation at human mendelian disease genes as a possible consequence of interference between advantageous and deleterious variants. eLife, 10:e69026, October 2021. ISSN 2050-084X. doi: 10.7554/eLife.69026. URL https://doi.org/10.7554/eLife.69026.

Daniel R Schrider. Background Selection Does Not Mimic the Patterns of Genetic Diversity Produced by Selective Sweeps. Genetics, 216(2):499–519, October 2020. ISSN 1943-2631. doi: 10.1534/genetics.120.303469. URL https://academic.oup.com/genetics/article/216/2/499/6066173.

Yassine Souilmi, M. Elise Lauterbur, Ray Tobler, Christian D. Huber, Angad S. Johar, Shayli Varasteh Moradi, Wayne A. Johnston, Nevan J. Krogan, Kirill Alexandrov, and David Enard. An ancient viral epidemic involving host coronavirus interacting genes more than 20,000 years ago in East Asia. Current Biology, 31(16):3504–3514.e9, August 2021. ISSN 09609822. doi: 10.1016/j.cub.2021.05.067. URL https://linkinghub.elsevier.com/retrieve/pii/S0960982221007946.

Daniel R. Schrider and Andrew D. Kern. Soft Sweeps Are the Dominant Mode of Adaptation in the Human Genome. Molecular Biology and Evolution, 34(8):1863–1877, August 2017. ISSN 0737-4038. doi: 10.1093/molbev/msx154. URL https://doi.org/10.1093/molbev/msx154.

Bernard Y Kim, Christian D Huber, and Kirk E Lohmueller. Inference of the Distribution of Selection Coefficients for New Nonsynonymous Mutations Using Large Samples. Genetics, 206(1):345–361, May 2017. ISSN 1943-2631. doi: 10.1534/genetics.116.197145. URL https://academic.oup.com/genetics/article/206/1/345/6064197.

Fumio Tajima. EVOLUTIONARY RELATIONSHIP OF DNA SEQUENCES IN FINITE POPULATIONS. Genetics, 105(2): 437–460, October 1983. ISSN 1943-2631. doi: 10.1093/genetics/105.2.437. URL https://doi.org/10.1093/genetics/105.2.437.

G. A. Watterson. On the number of segregating sites in genetical models without recombination. Theoretical Population Biology, 7(2):256–276, April 1975. ISSN 0040-5809. doi: 10.1016/0040-5809(75)90020-9. URL https://www.sciencedirect.com/science/article/pii/0040580975900209.

Y X Fu and W H Li. Statistical tests of neutrality of mutations. Genetics, 133(3):693–709, March 1993. ISSN 1943-2631. doi: 10.1093/genetics/133.3.693. URL https://doi.org/10.1093/genetics/133.3.693.

Justin C Fay and Chung-I Wu. Hitchhiking Under Positive Darwinian Selection. Genetics, 155(3):1405–1413, July 2000. ISSN 1943-2631. doi: 10.1093/genetics/155.3.1405. URL https://doi.org/10.1093/genetics/155.3.1405.

Kai Zeng, Yun-Xin Fu, Suhua Shi, and Chung-I Wu. Statistical Tests for Detecting Positive Selection by Utilizing High-Frequency Variants. Genetics, 174(3):1431–1439, November 2006. ISSN 1943-2631. doi: 10.1534/genetics.106.061432. URL https://doi.org/10.1534/genetics.106.061432.

Guillaume Achaz. Testing for Neutrality in Samples With Sequencing Errors. Genetics, 179(3):1409–1424, July 2008. ISSN 1943-2631. doi: 10.1534/genetics.107.082198. URL https://doi.org/10.1534/genetics.107.082198.

Guillaume Achaz. Frequency Spectrum Neutrality Tests: One for All and All for One. Genetics, 183(1):249–258, September 2009. ISSN 1943-2631. doi: 10.1534/genetics.109.104042. URL https://doi.org/10.1534/genetics.109.104042.

Anna Ferrer-Admetlla, Mason Liang, Thorfinn Korneliussen, and Rasmus Nielsen. On Detecting Incomplete Soft or Hard Selective Sweeps Using Haplotype Structure. Molecular Biology and Evolution, 31(5):1275–1291, May 2014. ISSN 0737-4038. doi: 10.1093/molbev/msu077. URL https://doi.org/10.1093/molbev/msu077.

Florencia Schlamp, Julian van der Made, Rebecca Stambler, Lewis Chesebrough, Adam R. Boyko, and Philipp W. Messer. Evaluating the performance of selection scans to detect selective sweeps in domestic dogs. Molecular Ecology, 25(1):342–356, January 2016. ISSN 1365-294X. doi: 10.1111/mec.13485. URL https://onlinelibrary.wiley.com/doi/10.1111/mec.13485.

John K Kelly. A Test of Neutrality Based on Interlocus Associations. Genetics, 146(3):1197–1206, July 1997. ISSN 1943-2631. doi: 10.1093/genetics/146.3.1197. URL https://doi.org/10.1093/genetics/146.3.1197.

Katherine M Siewert and Benjamin F Voight. BetaScan2: Standardized Statistics to Detect Balancing Selection Utilizing Substitution Data. Genome Biology and Evolution, 12(2):3873–3877, February 2020. ISSN 1759-6653. doi:10.1093/gbe/evaa013. URL https://doi.org/10.1093/gbe/evaa013.

Bárbara D Bitarello, Cesare de Filippo, João C Teixeira, Joshua M Schmidt, Philip Kleinert, Diogo Meyer, and Aida M Andrés. Signatures of Long-Term Balancing Selection in Human Genomes. Genome Biology and Evolution, 10(3):939–955, March 2018. ISSN 1759-6653. doi: 10.1093/gbe/evy054. URL https://doi.org/10.1093/gbe/evy054.

Leo Speidel, Marie Forest, Sinan Shi, and Simon R. Myers. A method for genome-wide genealogy estimation for thousands of samples. Nature Genetics, 51(9):1321–1329, September 2019. ISSN 1546-1718. doi: 10.1038/s41588-019-0484-x. URL https://www.nature.com/articles/s41588-019-0484-x. Publisher: Nature Publishing Group.

Thomas C. A. Smith, Peter F. Arndt, and Adam Eyre-Walker. Large scale variation in the rate of germ-line de novo mutation, base composition, divergence and diversity in humans. PLOS Genetics, 14(3):e1007254, March 2018. ISSN 1553-7404. doi:10.1371/journal.pgen.1007254. URL https://journals.plos.org/plosgenetics/article?id=10.1371/journal.pgen.1007254. Publisher: Public Library of Science.

Jacob A. Tennessen, Abigail W. Bigham Timothy D. O’Connor, Wenqing Fu, Eimear E. Kenny, Simon Gravel, Sean McGee, Ron Do, Xiaoming Liu, Goo Jun, Hyun Min Kang, Daniel Jordan, Suzanne M. Leal, Stacey Gabriel, Mark J. Rieder, Goncalo Abecasis, David Altshuler, Deborah A. Nickerson, Eric Boerwinkle, Shamil Sunyaev, Carlos D. Bustamante, Michael J. Bamshad, Joshua M. Akey, Broad GO, Seattle GO, and on behalf of the NHLBI Exome Sequencing Project. Evolution and Functional Impact of Rare Coding Variation from Deep Sequencing of Human Exomes. Science, 337 (6090):64–69, July 2012. ISSN 0036-8075, 1095-9203. doi: 10.1126/science.1219240. URL https://www.science.org/doi/10.1126/science.1219240.

Jeffrey R Adrion, Christopher B Cole, Noah Dukler, Jared G Galloway, Ariella L Gladstein, Graham Gower, Christopher C Kyriazis, Aaron P Ragsdale, Georgia Tsambos, Franz Baumdicker, Jedidiah Carlson, Reed A Cartwright, Arun Durvasula, Ilan Gronau, Bernard Y Kim, Patrick McKenzie, Philipp W Messer, Ekaterina Noskova, Diego Ortega-Del Vecchyo, Fernando Racimo, Travis J Struck, Simon Gravel, Ryan N Gutenkunst, Kirk E Lohmueller, Peter L Ralph, Daniel R Schrider, Adam Siepel, Jerome Kelleher, and Andrew D Kern. A community-maintained standard library of population genetic models. eLife, 9:e54967, June 2020. ISSN 2050-084X. doi: 10.7554/eLife.54967. URL https://elifesciences.org/articles/54967.

Benjamin C Haller, Peter L Ralph, and Philipp W Messer. SLiM 5: Eco-evolutionary Simulations Across Multiple Chromosomes and Full Genomes. Molecular Biology and Evolution, 43(1): msaf313, January 2026. ISSN 0737-4038, 1537-1719. doi: 10.1093/molbev/msaf313. URL https://academic.oup.com/mbe/article/doi/10.1093/molbev/msaf313/8342840.

